# Bayesian optimisation for yield in high-dimensional trait-space identifies crop ideotypes in Oil Seed Rape

**DOI:** 10.1101/2021.07.19.452946

**Authors:** Alexander Calderwood, Laura Siles, Peter J. Eastmond, Smita Kurup, Richard J. Morris

## Abstract

The improvement of crop yield has long been a major breeding target and is increasingly becoming a goal in many areas of plant research. Yield has been shown to be a complex trait, depending on multiple genes, plant architecture and plant-environment interactions. This complexity is frequently reduced by focussing on contributing factors to yield (yield traits). However, a quantitative understanding of the interplay between yield traits, and the effect of these relationships on yield is largely unexplored. Consequently, the extent to which crop varieties achieve their optimal morphology in a given environment and how this impacts on seed yield is unknown.

Here we use causal inference to model the hierarchically structured effects of 27 macro and micro yield traits on each other over the course of plant development, and on seed yield in Spring and Winter oilseed rape plants. We perform Bayesian optimisation on the modelled yield potential, identifying the morphology of ideotype plants which are expected to be higher yielding than the existing varieties in the studied panels. We find that existing Spring varieties occupy the optimal regions of trait-space, but that potentially high yielding strategies are unexplored in extant Winter varieties.

In addition to concrete recommendations for varietal improvement in oilseed rape, this work provides a novel, general methodological framework for the study of crop breeding as an optimisation problem.

## Introduction

Crop plants are often studied from the perspective of how to increase yield. However, yield is a complicated trait which can be decomposed into multiple yield components (yield traits) (Diepenbrock, 2000). For example, in oilseed rape (OSR), seed yield per plant can be expressed as seed size multiplied by the number of seeds. However, each of these primary yield traits can also be further factorised into secondary yield traits. For example, number of seeds can be expressed as seed number per pod x number of pods, meaning that seed yield can alternatively be expressed as seed size x seed number per pod x number of pods. Secondary traits can in turn be further decomposed into tertiary traits (and so on). These yield traits often exhibit complicated trait-trait relationships caused by physiological interactions.

Plants can be represented by points in “trait-space” – the space described by all possible combinations of yield trait values, in which each yield trait is a dimension. The trait-space is thus the domain of some (unknown) function which maps yield traits to yield potential. Plants exist on an (unknown) permissible manifold within trait space, defined by the constraints imposed by causal relationships between traits.

Crop improvement through genetic selection is a search for the permissible point in trait-space, which results in the highest yield potential. Sampling of trait space is achieved by iterating the two-steps of 1) making crosses between chosen parent plant varieties, and then 2) selecting lines within the progeny based on desired attributes for further evaluation. These steps are expensive, and so an efficient exploration method (providing maximum information from a given number of samples) is important. Two fundamentally different approaches to the search problem are possible.

In the first, “greedy” approach, crosses and selection are made using existing plants which perform better than their rivals. Selection is made either “directly” on the trait of interest (predominantly some measure of yield), or “indirectly” on other traits. Indirect traits for selection are identified, either as traits which have been observed to change in historic sets of cultivars exhibiting yield progress through breeding (Fischer, 2001), or as traits which are predicted through modelling approaches to associate with yield, but which are either easier to experimentally assay, or are more highly heritable (Bennett et al., 2017; Diepenbrock, 2000; Engqvist & Becker, 1993; Marjanović-Jeromela et al., 2007; Tariq et al., 2020).

However, depending on the (unknown) complexity of yield potential as a function of trait space, this greedy approach is potentially subject to inefficiencies (**see** **Figure 1**) and may also struggle to progress beyond potential local maxima in the yield surface. Crossing high-yielding parents without understanding why they are high-yielding may lead to the eventual perfection of a suboptimal yield strategy (finding a local maxima), alternatively the crossing of high-yielding parents with incompatible yield strategies, may lead to poor performing offspring (when a plateau, or yield minima exists between the parents in trait space).

**Figure 1:**
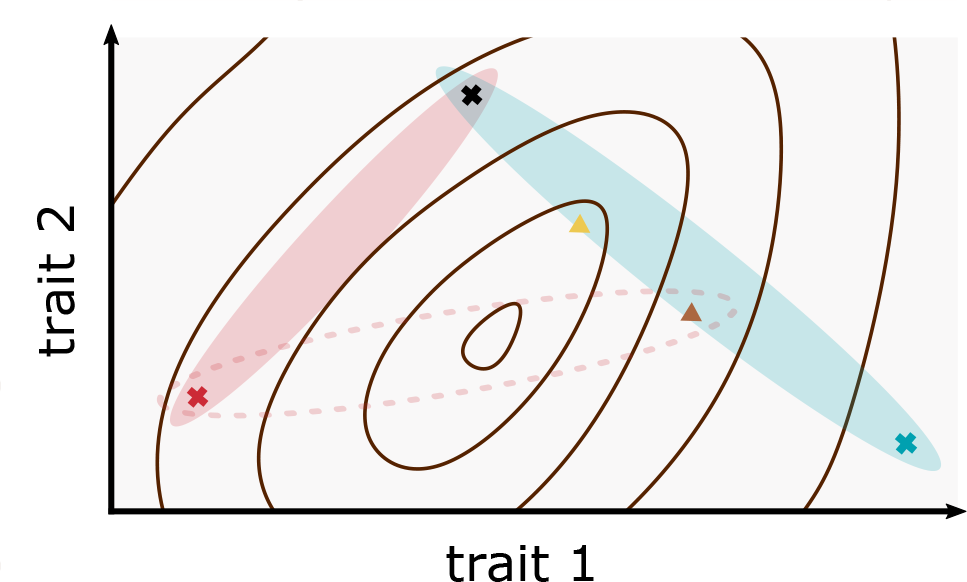
Identification of the yield surface facilitates selection of parent varieties for crosses, and selection among the progeny of a cross. A cartoon showing “trait-space”, with yield as a function of trait 1 and trait 2, indicated by contours with a maximum value in the middle of the plot. Plant varieties occupy points in the trait-yield space, marked as coloured crosses or triangles. Candidate parental lines are shown as crosses, offspring lines are shown as triangles. The trait values of potential offspring of a particular parental cross are abstracted as ovals in trait-space, reflecting that progeny phenotypes are not a linear combination of parent values, but are constrained by them. “Greedy” selection of parents selects the best yielding varieties to cross. Alternatively, “ideotype” selection may be done via an understanding of yield potential as a function of trait space. Either a qualitative understanding, (each parent contributes some “desirable” trait aspect), or a quantitative understanding (through a model of the yield surface in trait space). Even in this simple convex yield function example, greedy selection can be inefficient. In deciding between crossing either the red or blue candidate parent line with black parent, greedy selection favours crossing the red, as this yields higher than the blue parent. However, an understanding of the yield surface shows that progeny from a blue-black cross can be expected to include higher yielding varieties than from a red-black cross, as the blue oval covers a region of trait space with a higher maximum yield than the red oval. Knowledge of the yield surface is also useful in selecting which progeny of a cross to take forward. The yellow triangle plant is the highest yielding among progeny of the blue-black cross, but the suboptimal brown plant is worth retaining, as it can be crossed with the red plant to produce the best plants of all.

A second, hypothesis-driven approach, is to define (from biological expertise based on a physiological understanding) the trait values of the ideal hypothetical plant for maximum yield (an ideotype, Donald, 1968), which is then bred towards. Given a full description of the traits-to-yield function, the ideotype approach has the potential to be much more powerful than the “greedy” approach (**see** **Figure 1**). Given an understanding of how yield varies as a function of the yield traits, it is possible to select parent varieties with complementary traits to produce superior offspring, allowing convergence to the optimal yielding variety in fewer iterations, and avoiding local maxima. However, due to the complex causal relationships between traits, defining an ideotype in OSR based on biological expertise alone, without modelling is extremely difficult.

Causal trait-trait relationships may be based either on 1) definitional association (as described above), 2) developmental association (for example number of seeds is related to number of ovules because one develops into the other), 3) intra-plant competition for resources (for example pod number, seed number per pod, and seed size compete for seed filling resources, Diepenbrock, 2000), or 4) intra-plant signalling (for example seed derived signalling cascades trigger localised pod expansion (Pechan & Morgan, 1985; Ripoll et al., 2019).

Causal relationships between yield traits have been experimentally demonstrated through perturbation experiments (Iglesias & Miralles, 2014; Kirkegaard et al., 2018; Labra et al., 2017; N. Li et al., 2019; Pinet et al., 2015; Verdejo & Calderini, 2020; Wang et al., 2011; Zhang et al., 2020), or through careful studies of trait dynamics over time (Pechan & Morgan, 1985). Further, putative causal relationships have been identified between traits which are empirically associated (for example through correlation), and which can reasonably be expected to be developmentally related. For example, it has been demonstrated that 75% of the variation in pod number between plants can be attributed to differences in flower number (S. Li et al., 2020). These interactions can mean that a particular trait is beneficial for yield given one combination of other traits, but detrimental given another, for example, increased sink size may lead to increased yield given sufficient seed filling capacity, but otherwise be detrimental.

Trait-trait relationships can also result in changes in one trait causing changes in other traits which partially or completely nullify the intended effect on yield. For example, experimentally observed trade-offs between pod number, seed size, and seed number (Iglesias & Miralles, 2014; Kirkegaard et al., 2018; Labra et al., 2017; Pinet et al., 2015; Verdejo & Calderini, 2020; Wang et al., 2011; Zhang et al., 2020) make it difficult to intuitively predict whether increasing or decreasing each of these traits individually is expected to increase yield.

A further confounding factor in intuitive prediction is that many trait-trait relationships appear to be non-linear. Consequently, traits may have optimal values without being generally beneficial or detrimental. For example, experiments suggest that in general, yield in OSR is limited by seed filling capacity rather than by sink size (i.e. seed number and maximum size), but that experimental perturbation of traits can lead to a switch in this relationship (Zhang & Flottmann, 2018). The switch between these factors means that relationships between many traits (which encode aspects of either source production or sink size) and yield will be non-linear, so that they may affect it in some part of their value range, but not in others.

The complexity of a hierarchy of semantically nested and physiologically related yield traits can make identifying the occupiable region of trait-space difficult, and predicting the sensitivity of yield to trait location challenging. Mathematical modelling can help, both to understand the trait system given current data, and to propose the most informative new data to collect.

Observed correlations between traits can be partitioned into direct and indirect causal relationships via path analysis (Kozak et al., 2007; Marjanović-jeromela et al., 2008; Sabaghnia et al., 2010). In “sequential path analysis” a system of (*a priori*) known and permissible trait-trait relationships is defined, and the model of relationships which best explains the observed trait data given these constraints is inferred (reviewed Kozak & Azevedo, 2014).

Among modelling approaches to understand trait-trait relationships, sequential path analysis therefore offers a middle-ground between purely empirical modelling of correlations, and more principled physiological models of phenology (Deligios et al., 2013; Robertson & Lilley, 2016). Physiological models promise a complete and generalisable understanding of trait relationships under different environments, but require large amounts of detailed physiological data to parameterise.

To efficiently apply selective breeding after a desired trait state is identified, trait heritability must be confirmed and genetic markers underlying key traits identified. Sources of variation in a trait can be partitioned into genetic, environmental, and stochastic variability. Broad sense heritability is defined as the proportion of the variation in the trait attributable to genetic variation (Singh et al., 1993).

When causal trait-trait relationships are considered, sources of variation can alternatively be partitioned into the “direct” effects of genetics, environment, and intrinsic variability of the trait of interest, and the “indirect” effects that these factors have on this trait via their influence on the other traits which cause it. This partitions broad sense heritability into “direct heritability” (heritability of the trait, independent of other traits under selection), and “indirect heritability” (heritability caused by the heritability of the traits which affect it).

Quantifying the magnitude of the direct and indirect heritability of traits is important for physically producing a multi-trait ideotype plant, as otherwise, trait-trait relationships between co-selected traits may restrict the response to selection (for example if one desired trait causes another trait which is to be selected against).

GWAS studies may be used to identify the genes underlying genetic variation in candidate yield traits. Trait-trait relationships also have complicating effects on these studies, as they may cause genetic effects to be attributed to misleading traits. Cross-phenotype associations may be caused by biological pleiotropy, mediated pleiotropy, or by genetic linkage or poor experimental design and controls (Solovieff et al., 2013). Mediated pleiotropy occurs when phenotypes are causally related, and so any genetic loci associated with the first (causal) phenotype is also statistically associated with the second. In OSR, variants at several genetic loci have been associated with multiple yield traits. For example (N. Li et al., 2019; P. Yang et al., 2012) identify overlapping QTL for the causally related traits silique length, and seed weight (N. Li et al., 2019; Pechan & Morgan, 1985), and (S. Li et al., 2020) identify genes known to regulate flower number in a GWAS testing for genetic association to silique number.

Distinguishing biological pleiotropy from mediated pleiotropy is valuable to avoid the misleading identification of genes as being directly involved in a trait of interest but which in fact act indirectly via a different trait. One can distinguish biological pleiotropy from mediated pleiotropy by controlling for the traits which cause each trait of interest (Solovieff et al., 2013; Vansteelandt et al., 2009).

Here, we explore the consequences of causal trait-trait relationships for crop improvement through their effect on predictions for yield improvement, trait heritability and genetic association studies. OSR is an excellent system for studying trait-trait relationships due to its complicated and plastic developmental morphology, and overlap between the periods of branch growth, flowering, pod setting, and seed filling (Diepenbrock, 2000; Iglesias & Miralles, 2014; Kirkegaard et al., 2018; Tayo & Morgan, 1975). Studying the consequences of causal trait-trait relationships is timely as, with phenomics platforms becoming more widely available, it is becoming increasingly possible to observe large numbers of potentially related traits, and so to account for these relationships statistically.

We measure macrotraits (measured at the whole plant level) and microtraits (comprising ovule area and number, gynoecia, ovary and style length) to comprehensively study traits previously associated with yield, as well as the less studied female reproductive traits (Chen et al., 2014; Jiao et al., 2021; Özer & Oral, 1999; Tariq et al., 2020; Wang et al., 2011; Y. Yang et al., 2017).

Through sequential path analysis we identify and model trait-trait relationships and the trait-yield function. Consequently, we identify which single traits are the best candidates for modification to alter yield potential in Spring and Winter OSR, accounting for compensatory or exacerbating changes in other traits.

We apply Bayesian optimisation of yield as a function of plant morphology to identify the trait combinations which should be prioritised for exploration through selective breeding to advance OSR improvement most efficiently. These points define OSR crop ideotype morphology – the hypothetical optimal combinations of yield-trait values that are expected to produce the greatest possible seed yields under our growing conditions, when constrained by the observed trait-trait relationships to occupy the feasible region of trait space, and explore currently unoccupied regions of it.

We use the inferred trait-trait models to highlight the impact of causal trait relationships on estimates of heritability, and gene association studies. Using numerical simulation we demonstrate that such interactions make the inference of genetic associations challenging to identify statistically, but that by conditioning on causal traits, we can not only distinguish biological pleiotropy from mediated pleiotropy, but also improve the power to detect genetic variants associated with a trait of interest. Using several examples we show that the approaches described here provide a novel, general methodological framework for the study of crop breeding as an optimisation problem.

## Results

### Trait-trait correlations are caused by direct and indirect relationship structure

OSR exhibits complex morphology with many traits which might be expected to affect seed yield. We measured 27 traits in Spring and Winter OSR to examine relationships between them and with seed yield.

Spring and Winter OSR morphology differs (Chen et al., 2014), and studies conducted in Spring type and Winter type OSR have reached different conclusions about key features in development of crop yield (Diepenbrock, 2000; Siles et al., 2020), so we analysed Spring and Winter OSR varieties separately.

Many traits are highly correlated with one another (**Figure 2**), however the causal structure underlying the complex relationships between the traits is not obvious from this correlation matrix. In order to understand the identities and strengths of these trait-trait relationships, and so be able to predict the effect on seed yield of perturbing traits (for example through selective breeding), we used sequential path analysis (reviewed by Kozak & Azevedo, 2014) to fit a model which best explains how these traits affect each other and seed yield (see Methods).

**Figure 2:**
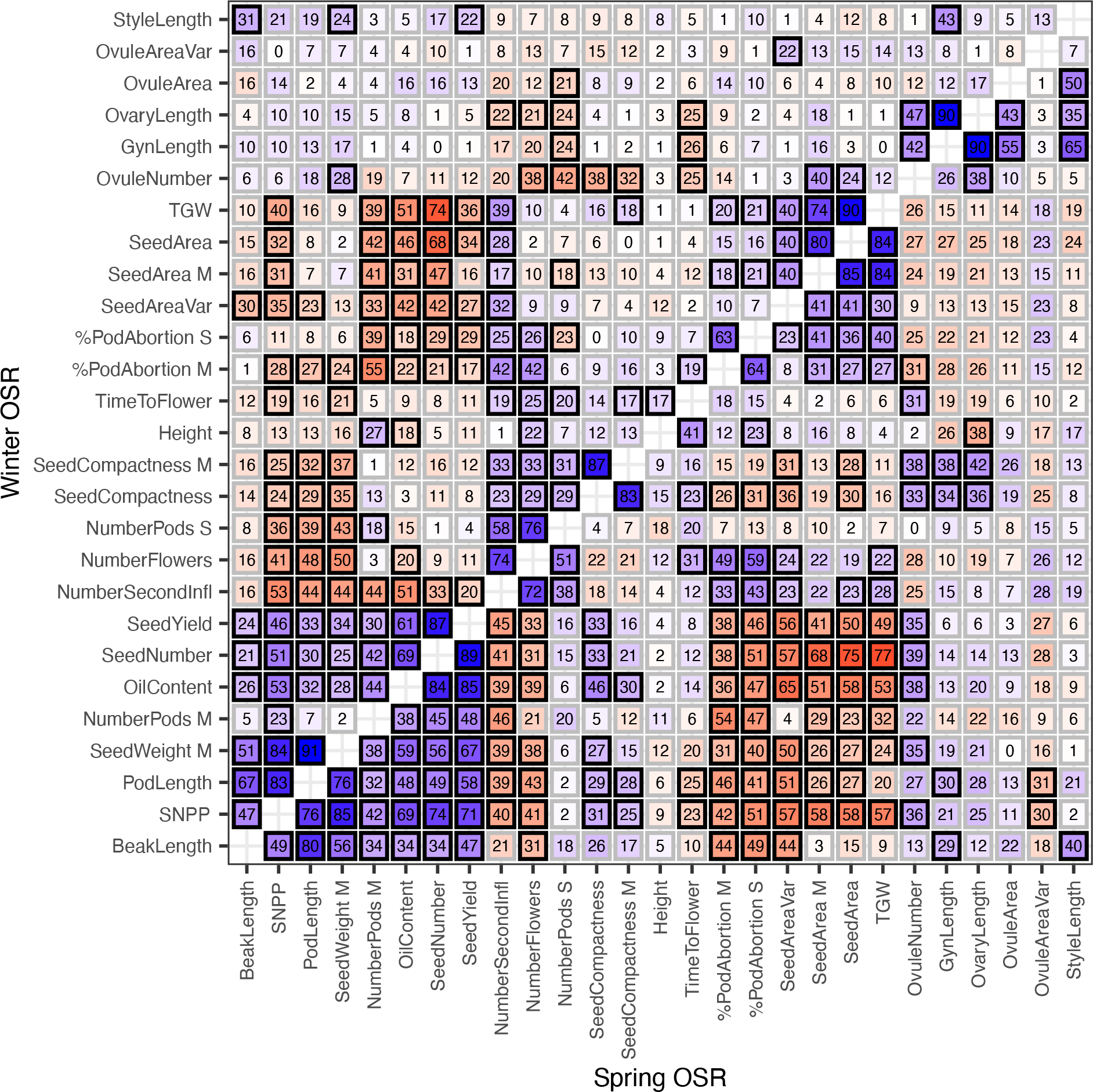
Correlation between traits in Spring and Winter OSR accessions. Spearman’s correlation coefficients between traits for Winter (above diagonal) and Spring (below diagonal) OSR accessions. Blue shows positive correlation, red shows negative. Reported values are absolute values of coefficients, multiplied by 100. A black border indicates statistically significant, non-zero correlation (t-test), using Benjamini-Hochberg adjusted p-value with significance level 0.05.

**Figure 3** shows the directed acyclic graph (DAG) structure of the inferred model for traits and seed yield (SeedYield) in Spring and Winter OSR. Traits which are connected directly were inferred as having a direct causal relationship. For example, in Spring OSR, the number of flowers (NumberFlowers) directly affects the number of pods (NumberPods S). Conversely, nodes which are indirectly linked via intermediate nodes have an indirect causal relationship, mediated by the intermediate traits. For example, in Spring OSR the number of flowers (NumberFlowers) affects seed yield indirectly: the number of flowers (NumberFlowers) affects number of pods on secondary branches (NumberPods S), and the number of pods then affects seed yield. Other indirect causal paths between the number of flowers, and seed yield can also be seen, for example via the number of pods on the main inflorescence (NumberPods M).

**Figure 3:**
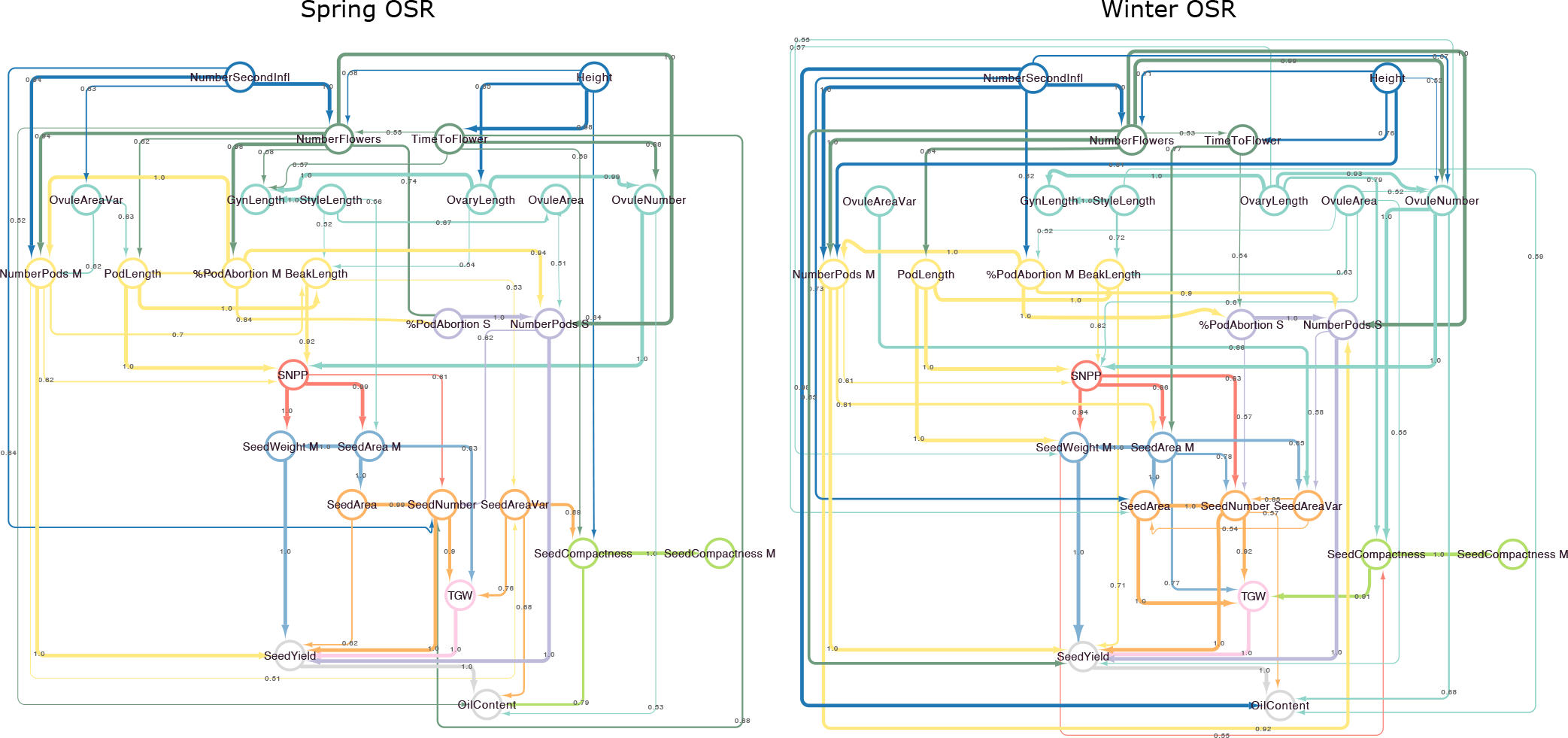
Graphical models for inferred relationships between traits in Spring and Winter OSR. Observed traits are shown as nodes, edges link traits which are inferred to be directly associated, without mediation by other observed traits. Edge directionality is from the inferred causal trait (parent trait) to the caused trait (child trait). Node position and colour corresponds to the predefined hierarchy of traits (see methods). Edge colour corresponds to the source node colour. Edge width, and annotation show the estimated probability of the existence of that trait-trait relationship.

Many relationships between the measured traits and seed yield are indirect, being attributable to the effect of the trait on other intermediate traits, which then themselves affect seed yield either directly or indirectly.

The inferred model graphs are consistent with well-known relationships, for example in both Spring and Winter OSR, the number of flowers (NumberFlowers) is directly caused by the number of secondary inflorescences (NumberSecondInfl) and the number of pods on the main, and secondary branches (NumberPods M and NumberPods S) are consequences of the number of flowers and pod abortion (%PodAbortion).

The inferred DAGs also identify differences in trait relationships between ecotype groups. For example, in Spring OSR, flowering time (TimeToFlower) is predicted to affect the seed number per pod (SNPP, via OvuleNumber), but in Winter OSR accessions it is not, presumably because in these slower flowering varieties, TimeToFlower imposes less of a constraint upon the seed filling potential of the plant.

### Trait-trait relationships buffer effects of trait modification on seed yield

Although the inferred DAG structures identify the paths by which traits affect each other and ultimately seed yield, their complexity makes it difficult to read the effect that any trait modification is expected to have. We therefore used the inferred graph structures to model the effect that changing yield traits can be expected to have on seed yield in Spring and Winter OSR (see methods).

**Figure 4** shows predicted seed yield for Spring and Winter OSR as each trait is varied individually, whilst holding upstream traits to their observed values, but allowing downstream traits, and seed yield to vary due to the altered trait.

**Figure 4:**
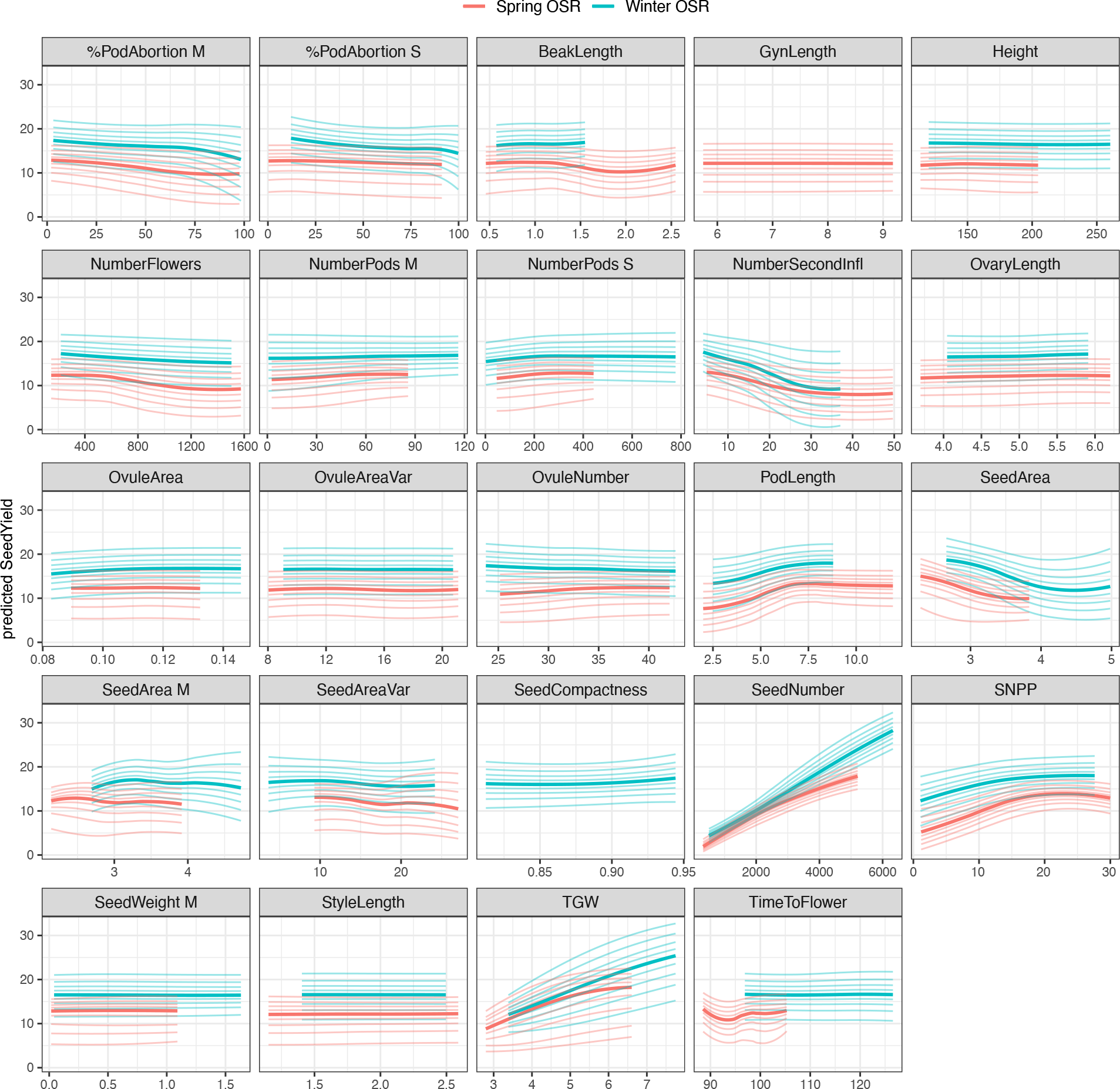
For many yield traits there is insufficient evidence to conclude that their individual modification will generally alter yield. Trait-trait models were used to predict the effect of altering yield trait values in Spring and Winter OSR assuming directly linked traits are linked by causal relationships. Each facet shows the predicted effect of changing one yield trait on seed yield between the minimum and maximum values observed for that trait. Traits downstream of the altered trait in the model DAGs were predicted, and so allowed to vary in consequence of the changes. All traits upstream of, (or disconnected from) the varied trait were held to their observed values. Only plots for traits which are upstream of seed yield in the modelled DAGs (and which therefore may affect it) are shown. Yield values were predicted for all varieties in each panel. Median predicted seed yield values are shown in heavy line. Predicted seed yield quantiles between 10% and 90% at 10% intervals are plotted with light lines. Uncertainty is a consequence of both uncertainty in the modelled relationships, and variation between the varieties in each panel. Spring OSR is shown in red, Winter OSR in blue.

Strikingly, we see little clear evidence that individually modifying many traits will affect seed yield. Traits which are not predicted to affect Seed Yield can be divided into two classes.

One class contains traits which are not correlated with Seed Yield, even though causal paths exist between them in the model DAG structure. For example, the number of flowers (NumberFlowers) in Winter OSR (see **Figure 2**, **Figure 4**). The lack of effect that members of this group have on yield is due to compensatory changes in other traits. For example, although a greater number of flowers is predicted to result in a greater number of pods, these pods are expected to be shorter, and each contain fewer seeds, resulting in little predicted effect on seed yield (**Figure 5**).

**Figure 5:**
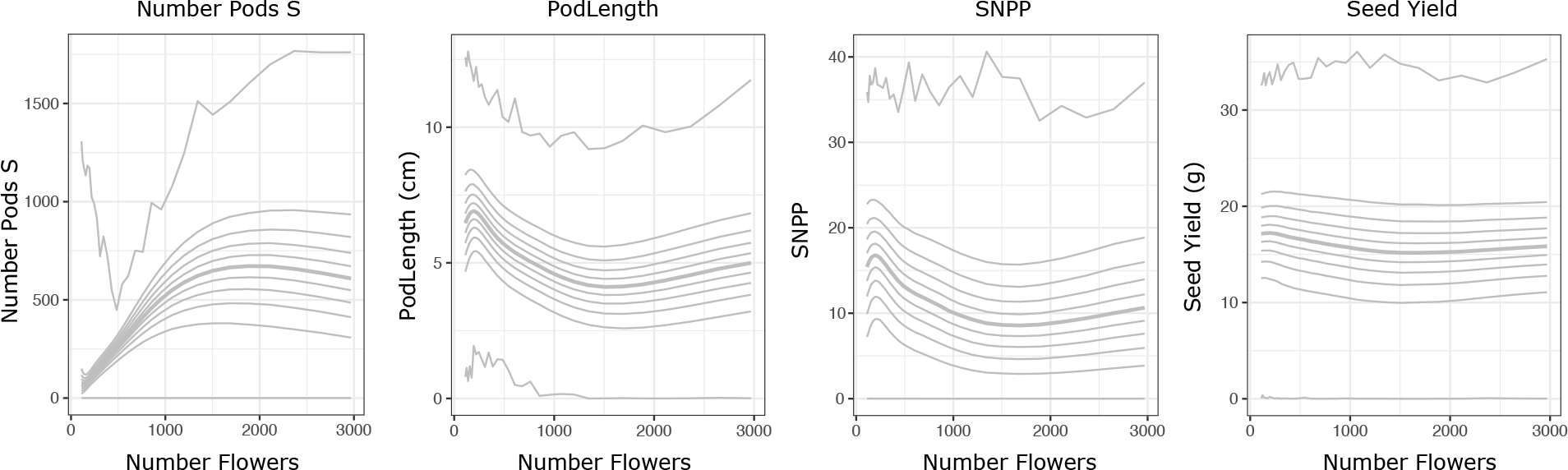
Compensatory interactions between yield traits buffer the effects of many yield traits. The predicted effects in Winter OSR of modifying the number of flowers on the number of pods on secondary branches (Number Pods S), average pod length (PodLength), seed number per pod on the main branch (SNPP), and seed yield per plant (Seed Yield). Median value, and 10% quantiles in the response trait predictions are shown. Although increasing the number of flowers is expected to result in the production of more pods (up to a point), compensatory changes in pod length and seed number per pod result in an expected slight reduction in final seed yield.

In the second class are traits which are highly correlated with yield, but whose modification is not expected to affect it. This is because these traits and yield have a common causal factor. For example, silique beak length (BeakLength) in Winter OSR: the main source of the association between beak length and seed yield is through pod length, which causing both yield and beak length. Consequently, modifying beak length is expected to have little impact on the other observed traits, and is not expected to effect seed yield, whereas modifying pod length is expected to affect several traits, including beak length and seed yield, (see **Supplemental Figure 1 a, b**).

**Figure 4** shows that the traits whose alteration are most confidently predicted to modify seed yield are the number of secondary inflorescences (NumberSecondInfl), pod length (PodLength), total seed number (SeedNumber), seeds per pod (SNPP) and thousand grain weight (TGW) and seed area.

Interestingly, increasing total seed number (SeedNumber) is expected to lead to increased seed yield, despite identification of the well-known negative relationship between seed number and TGW (**Supplemental Figure 1c,** Labra et al., 2017). This suggests that the trade-off between TGW and seed number is such that the production of many seeds is favoured, despite their reduced size. Note that although increasing TGW is also expected to increase seed yield, this is under the assumption that seed number (as an upstream node of TGW in the causal network) is held constant and so does not take account of the relationship between them.

Surprisingly similar seed yield responses to trait variation are predicted between Spring and Winter OSR varieties, although there are quantitative differences in response or optimal value for some traits. Notably, a small number of secondary inflorescences (NumberSecondInfl) appears beneficial in both types. This was expected in Winter OSR (Diepenbrock, 2000), but previous work has suggested a large number of branches to be beneficial in Spring OSR (Aytaç & Kınacı, 2009).

We see many examples of non-linear effects in the expected response of seed yield to perturbation of yield trait values. Therefore whether yield is expected to be improved through a trait’s modification in a particular variety depends on the existing value of that trait. For example, although at low values, a bigger seed number per pod on the main inflorescence (SNPP) is expected to increase seed yield, increasing the number continually is not expected to lead to significant further seed yield improvements in either Spring or Winter OSR (**Figure 4**), presumably unless buffering changes in other traits above this value can also be controlled.

It is therefore not clear that recommendations based on linear models which suggest that individual traits should be “higher” or “lower” are general across varieties, or can be considered in isolation from other traits. The exceptions to this are total seed number (SeedNumber) and thousand grain weight (TGW), which do not exhibit saturation behaviour in the expected yield response within the observed trait range (at least in Winter OSR).

### Trait-trait relationships reduce maximum possible seed yield and alter identity of OSR ideotypes

Having identified traits which individually are expected to affect seed yield, and observed non-linear, compensatory effects among them, we were interested to identify crop ideotypes in terms of multiple traits simultaneously. To identify ideotype plants under our experimental conditions, we performed Bayesian optimisation of trait values to maximise the expected improvement (EI) in seed yield (see Methods). Optimisation of EI allows identification of points which, given prediction uncertainty, have the greatest probability of generating a higher seed yield than the maximum among the observed plants. Hence these points in trait space represent proposed crop ideotypes; hypothetical plants which have the best chance of yielding better than the observed plants, but which are constrained to be somewhat distant (in trait space) from the existing, experimentally observed plants where the seed yield is already known, and so prediction uncertainty is relatively small.

To reduce the dimensionality of the trait space considered, we used only traits which directly affect seed yield (that is the traits which connect to it directly in **Figure 3**). Although as shown in **Figure 4**, modification of some indirectly acting traits is also expected to affect seed yield, their action is only via directly connected traits. Consequently, if the traits directly connected to seed yield are controlled, then all other traits are irrelevant to seed yield. It is only necessary to consider indirectly connected traits if the desired values of directly acting traits cannot be achieved without their modification (for example, because a more extreme value is required than is present in the pre-breeding material, see **Supplemental Figure 2**).

If the empirically observed relationships between the yield traits are ignored, then unsurprisingly, optimisation finds many ideotype plants that are expected to have higher seed yield than the experimentally observed plants in both the Spring and Winter OSR panels (see **Supplemental Figure 3**). Predominantly the proposed ideotypes produce very large numbers of very, or moderately heavy seeds, and a broad range of values for the other yield traits.

However, this relaxation of the observed correlations among traits is unlikely to be reasonable as previous experimental work indicates that they are due to intra-plant competition for seed filling resources (Iglesias & Miralles, 2014; Kirkegaard et al., 2018; Labra et al., 2017; Pinet et al., 2015; Verdejo & Calderini, 2020; Wang et al., 2011; Zhang et al., 2020).

**Figure 6** shows the observed yield of experimental plants, and the expected seed yields of ideotype plants when empirically observed relationships between yield traits are respected as additional constraints during the optimisation of EI (see methods).

**Figure 6:**
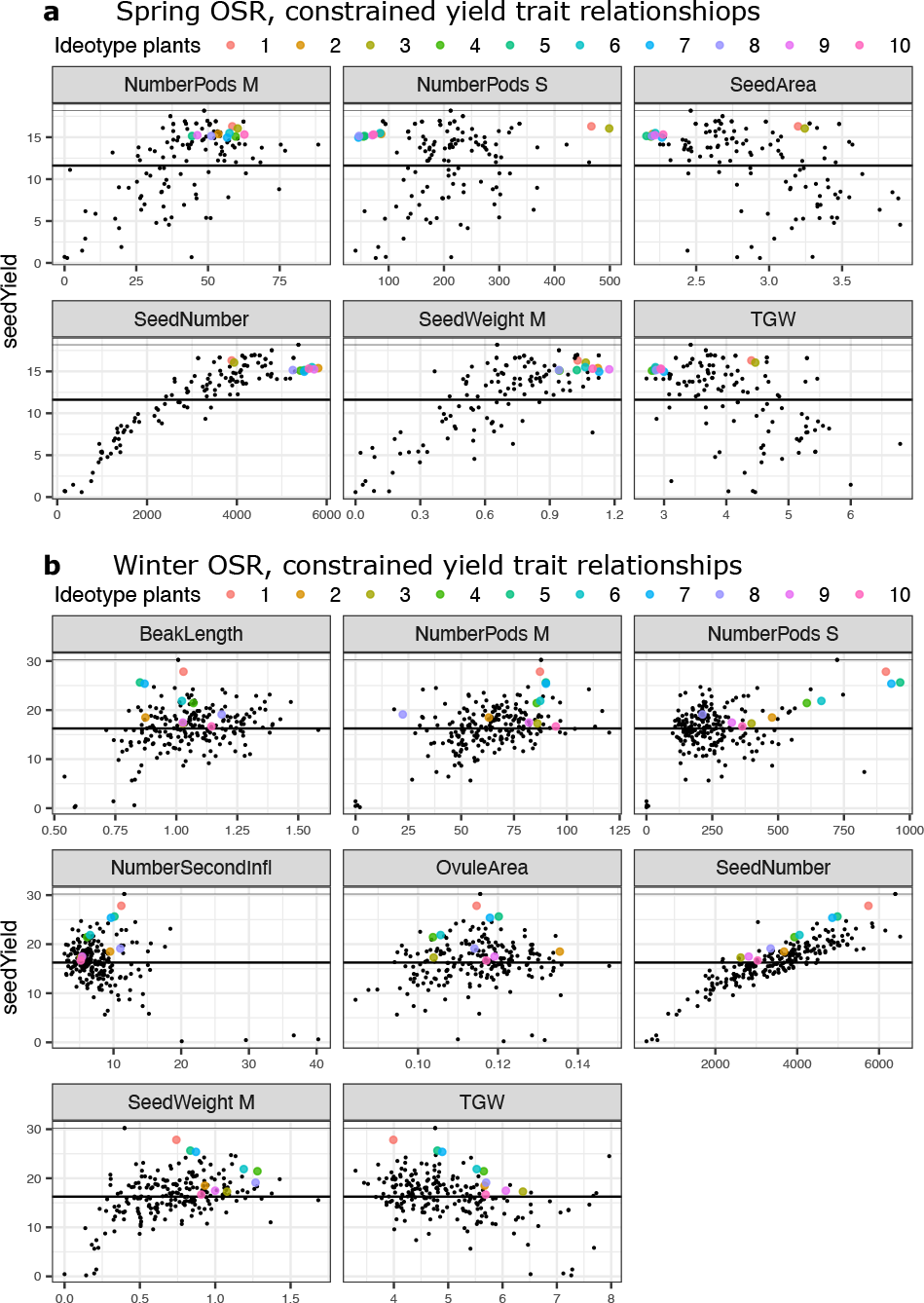
Model identified optimal crop ideotypes. Facets show traits which affect seed yield directly in the **a)** Spring and **b)** Winter OSR panels. Coloured points show mean predicted seed yield for identified hypothetical plants with the indicated yield trait values. Their order reflects their probability of having greater seed yield than the best experimentally observed plant, with 1 having the best chance, and 10 the worst of the calculated ideotypes. These points are identified through Bayesian optimisation as the unobserved points with the best chance of exceeding the maximum observed seed yield in each panel. Black points show seed yield and yield trait values for experimentally observed plants. The thin black line marks the maximum seed yield observed amongst the observed plants, the thick black line shows mean seed yield among the observed plants.

In Spring OSR, many observed plants are higher yielding than the expected values of any hypothetical plants. This indicates that when yield trait relationships are constrained to follow their observed relationships, the optimal regions of trait space are already well explored by existing plants, at least when grown under these environmental conditions.

The high-yield region is broadly defined by the production of a large number of seeds (SeedNumber > 3000) relative to the other plants in the panel, and a large proportion of productivity via the primary inflorescence relative to what is usual in Spring OSR. Seeds produced on the main inflorescence are relatively heavy (SeedWeight M > 0.6g), but the weight of seeds produced over all inflorescences (TGW) of high performing plants does not appear to be tightly constrained and covers most of the range observed among all Spring OSR plants. The number of pods on the main inflorescence is fairly restricted, to slightly above the average, ranging from approximately 35 – 60. The number of pods on other branches (NumberPods S), and seed area (SeedArea) are more variable, but on average slightly lower than in the general Spring OSR population.

A focus on production via the main inflorescence appears to be required for good performance. The most promising strategies for hypothetical ideotypes are to vary the approach to secondary branches (NumberPods S), whilst maintaining high production on the primary inflorescence. In contrast to the observed plants which have a middling number of pods on secondary branches (150 – 350), ideotype plants 1 and 3, have and extremely large number of pods on secondary branches (approximately 500), and produce slightly fewer, slightly larger seeds than the observed high performing lines. The other ideotype plants have a low number of pods on secondary inflorescences (< 100) and produce an overall larger number of smaller seeds than is usual among the experimentally observed plants.

Conversely, among Winter OSR, only one observed plant yields higher than the mean expected seed yield of the best hypothetical point. This suggests that the optimal yield trait region under these conditions may not have been fully explored by varieties in the Winter OSR panel.

In Winter OSR increasing seed number leads to increased seed yield, and the best ideotypes produce the most total seeds (seedNumber) of smaller size (TGW). Whilst the identified ideotypes respect the trade-off between seed number and TGW, the most promising ideotype plants occupy the upper edge of the cloud of experimentally observed plants in both these metrics. This means that they are expected to be able to produce heavier seeds than observed plants which produce the same number of seeds, and more seeds than plants which produce the same TGW.

To allow this, they have high values for traits associated with increased photosynthetic capacity relative to the Winter OSR average. Among the three highest yielding ideotype plants, the number of secondary inflorescences is between 10 and 12, (compared to an average of 8 among Winter OSR), and more than 850 pods on secondary branches are proposed, further increasing the photosynthetic area. This strategy is similar to that of the single best performing observed plant. Ideotype production on the main inflorescence is suggested to be similar to that observed among the experimentally high yielding plants. The best ideotypes have approximately 90 pods on the main inflorescence, and produce 0.75 – 0.9g of seed per 10 pods on the main stem (seedWeight M).

### Trait-trait relationships exaggerate trait heritability estimates

When choosing traits for genetic selection, it is important to ascertain that genetically controlled variation in the trait exists within the available breeding material. However, we find that many of the measured traits can be well predicted from observations of their parent traits **(see Suplemental** **Figure 4**). This implies that a large part of the observed variation in these traits may be caused indirectly by variation in their parent traits, rather than by direct variation in the trait of interest.

As shown in **Supplemental Table 1**, the majority of measured traits exhibit evidence for substantial “total” broad sense heritability. However, when variation caused by parent traits is accounted for, estimates of “direct” heritability can substantially decrease. For example, heritability for pod abortion on the main branch (%abortion M) drops by a factor of 3x when variation in its parent traits (number of flowers and number of secondary branches) are accounted for.

This indicates that when attempting to produce a plant defined by multiple traits, care must be taken in trait selection, as there is less potential for their independent modification than might be calculated naively.

### Trait-trait relationships mask genetic association unless controlled for

When traits can cause each other, a gene may have an indirect causal link to the (child) trait of interest, mediated by its direct effect on intermediate (parent) traits (see **Supplemental Figure 5a**). This is called mediated pleiotropy.

One method to distinguish genes acting directly upon a trait from genes acting indirectly (mediated by other traits) is to correct for the association between causally linked traits using the residuals of modelled trait-trait relationships (Vansteelandt et al., 2009). This eliminates association between the trait of interest and the parent trait(s), and therefore breaks the causal path between trait of interest, and any SNPs which affect it through the parent trait(s).

Under simulation, we find that when variation in the child trait is predominantly caused by variation in the parent trait, then unless the correction is applied, SNPs which act on the parent are preferentially identified over SNPs which act directly on the child trait **Supplemental Figure 5b**). Correction for trait-trait relationships also increases the power to detect causal SNPs (**Supplemental Figure 5 b**). Without correction, the child trait is effectively more genetically complex, being caused not only by the SNPs which affect it directly, but also by all the SNPs which effect the parent trait(s).

To check whether evidence of these behaviours can be seen in real data, we conducted GWAS analysis on traits measured in OSR panel, with and without correction for the effects of trait-trait relationships. **Figure 7** shows GWAS results for number of pods on the main inflorescence, and seed oil content.

**Figure 7:**
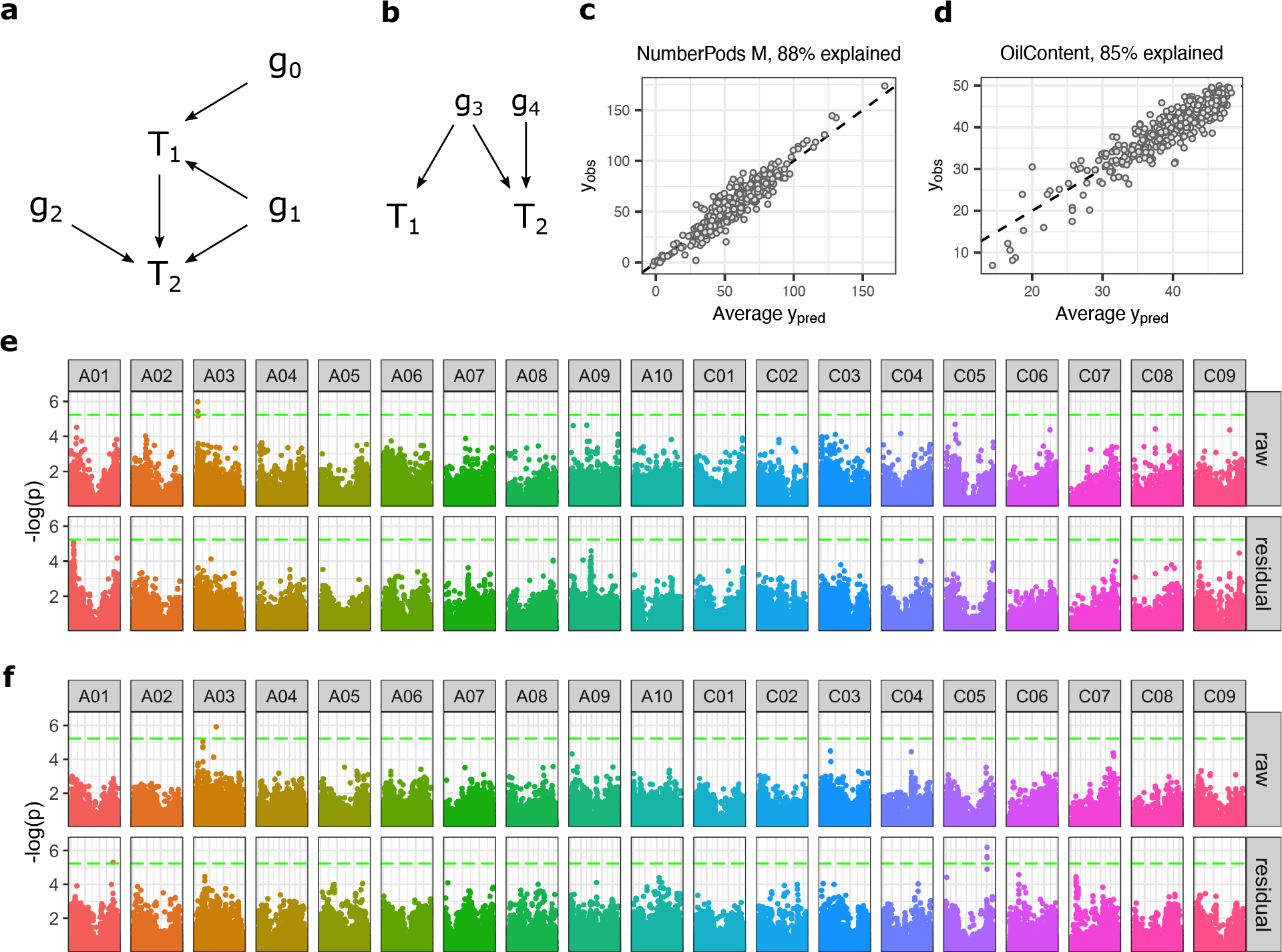
Accounting for trait-trait relationships in real data, can identify SNPs which likely act indirectly, and identify SNP associates which are missed when no correction for is applied. **a)** Causal diagram showing relationships between traits (T) and genes (g). If a causal relationship exists between traits, genes may affect a trait indirectly via another trait (g_0_, mediated pleiotropy), directly (g_2_), or via both of these mechanisms (g_1_). **b)** Trait-trait relationships may be erroneously inferred due to a confounding latent variable, for example a pleiotropic gene which causes both traits to be correlated (g_3_). **c)** Variation in the number of seed pods on the main inflorescence (NumberPods M), and **d)** seed oil content (OilContent) can be well explained by variation in their parent traits. Manhatten plots for SNPs associated to **e)** number of pods on the main inflorescence (NumberPods M), and **f)** seed oil content (OilContent). Association is tested either the raw trait (raw), or after correction for parent trait values (residual). Statistical significance is shown with Bonferroni corrected FWER = 0.1. Considering NumberPods M, a significant peak on chromosome A03 in the raw plots disappears in the residual plots. Assuming a causal relationship between the parent traits and NumberPods M, then this peak is likely caused by mediated pleiotropy. However, it is unclear whether this entirely or partly explains the association (g_0_, or g_1_ in **a**). If no causal link exists between the traits, then this correction can result in falsely discarding g_3_ type genes (see **b**). Considering OilContent, significant peaks appear on C05 and C07 in the residual plot, which are not present in the raw plot. If a causal link between the traits truly exists, this may be because accounting for variance in the parent traits increases the power to detect SNPs which affect the trait of interest (g_2_). If instead both traits have common genetic causes this can still result in increased power to detect genes affecting oil content by correcting for the variance introduced by these genes (by removing the effect of g_3_, g_4_ is more easily detected in **b**).

**Figure 7e** shows a statistically significant peak on chromosome A03 associated with the observed number of pods on the main inflorescence (see **Supplemental Data Set 1**). However, this peak (which is visible when using the “raw” data) disappears when using the parent trait corrected model residual data. Given casual links between the parent and child traits, then disappearance of a peak in the raw observation data may be because the gene under the peak acts on the trait of interest directly via mediated pleiotropy (as g_0_ in **Figure 7f** **a**).

This correction does increase the potential for falsely eliminating genes from consideration if genes are truly acting pleiotropically “directly” (g_1_ in **a**). If there is not truly a direct causal link between the traits, pleiotropic gene action may underly the association between the traits which leads to their inferred linkage in the model of trait relationships (**Figure 7b**, g_3_).

**Figure 7f** shows SNP association with seed oil content. A statistically significant peak on chromosome C05 appears after the trait relationship correction is applied **(Supplemental Data Set 2).** The detection of this peak using residual correction is due to increased power to detect SNPs after parent trait action is accounted for.

This increased power may be due to direct causal interaction between traits (in which case the gene under the appearing peak is analogous to g_2_ in **Figure 7a**), or because the inferred association between parent and child traits is actually caused by underlying genes which effect both traits (**Figure 7b**). The gene(s) affecting both traits may either be truly pleiotropic genes, or correlated genes (either through linkage disequilibria, or through joint selection). Variability in oil content caused by variation in these genes then masks the effect of the gene under the C05 peak (similar to g_4_) until their effects are indirectly controlled for.

In this example, it is likely that both mechanisms are relevant. As well as a plausible causal link between the traits themselves, seed yield and oil content are under joint selection, (both are positive traits for OSR breeding programmes), and so genes affecting these traits are likely to be correlated in the population.

## Discussion

Here, we have analysed the underexplored consequences of trait-trait relationships, finding that they alter key yield traits and ideotypes, heritability estimates, and gene-trait association studies.

We inferred the trait network structures which are most parsimoniously able to explain the empirically observed trait-trait relationships, consistent with prior beliefs about trait relationships. Establishing these network structures in OSR is important to understand how traits can potentially affect seed yield, but also how traits can affect each other. Highly connected networks were inferred for both Spring and Winter OSR. This supports the view many yield traits depend on, and affect each other.

Whilst our model allows for complex interactions to be captured, it does have limitations. We modelled trait-trait relationships as a directed acyclic graph (DAG). Therefore, feedback loops between traits cannot be accounted for in the model, although they are expected to exist in the plant (N. Li et al., 2019; Pechan & Morgan, 1985). Similarly, the trait-trait relationship networks are expected to change over development (Kirkegaard et al., 2018; Verdejo & Calderini, 2020; Wang et al., 2011; Zhang & Flottmann, 2018), and environmental conditions (**Supplemental Figure 6**). As functional trait data becomes available, it will allow increasingly sophisticated modelling of feedback between traits and dynamic network structures via dynamic Bayesian network approaches.

The inferred trait models were used to predict the impact of changing each trait on seed yield in Spring and Winter OSR, accounting for compensatory or exacerbating changes in other traits. The traits whose modification are most confidently predicted to lead to variation in seed yield when averaged across all of the studied varieties are the number of secondary inflorescences (NumberSecondInfl), pod length (PodLength), total seed number (SeedNumber), seeds per pod (SNPP) and thousand grain weight (TGW) and seed area (SeedArea), suggesting that these traits provide some of the most important levers for the improvement of yield.

Given the highly connected DAGs identified, it is surprising that for most traits, we cannot confidently predict that perturbation through selective breeding (at least within the range of observed values) will lead to an improvement in seed yield. This partly reflects inherent uncertainty in the modelled relationships between traits, (which accumulates with each additional indirect step between the perturbed trait and seed yield), but also (as exemplified by flower number) reflects the many compensatory and buffering relationships between traits in the developmentally plastic OSR plant structure. This is in contrast to results in wheat, which has been found to be relatively developmentally determinate (Asseng et al., 2017).

Differences between the predicted effect of traits in the sequential path model versus correlation analysis occur when (under the model), confounding trait-trait relationships cause the correlation. This can mean that traits which are correlated with seed yield may not be expected to have a strong causal effect on it. For example, the correlation between beak length and seed yield can largely be explained by the variables which cause them both.

Conversely, trait-trait relationships can also obscure, or invert correlations between traits and yield. Analysis of the structure of the trait-trait relationship DAGs allows identification of the compensatory relationships between traits which prevent a particular modification affecting yield, and so the identification of which trait-trait relationships would have to be broken to allow the trait to affect yield. Consider TGW, which has been suggested not to be used as a selection criteria, as it is not correlated with seed yield (Bennett et al., 2017). In our dataset, TGW is negatively correlated to seed yield (see **Figure 2**), but (through modelling) we find that it is expected that increasing TGW directly will increase yield. This apparent contradiction is due to the strong negative causal relationship from seed number to TGW (**Figure 2**) – small seed number leads to both low seed yield and large seed size, rather than large seeds necessarily leading to low seed yield. When this trait-trait relationship is accounted for, increasing TGW (whilst holding seed number fixed) is expected to lead to increased yield (see **Figure 4**), in spite of the negative correlation between TGW and seed yield.

Under these experimental conditions, we find that the qualitative relationships between traits and seed yield are similar in Spring and Winter OSR. Differences are largely in the optimal magnitude of traits, rather than in which traits are most important, or the sign of their effect on seed yield.

Among the traits which are expected to affect seed yield, we see many non-linear, saturation type curves. This suggests that the correlation coefficients, or linear models widely used in trait modelling may be misleading in identifying a particular secondary yield trait for selection for or against to improve seed yield. Instead for the majority of traits, an optimal region exists within the range of values already in available material. For most of the observed traits, we predict little benefit in more extreme trait values than are already present in the studied panels.

Identification of the individual traits whose modification is expected to alter seed yield provides a good way to identify “important traits” for yield modification, however it fails to identify which combination of multiple trait values is required for a high-performance variety as it ignores potential interactions between trait modifications.

A “trait space” describes the values that the observed traits can take. In this view, OSR plants exist on a manifold within trait space, defined by the constraints of relationships between traits. Rather than identifying the most important individual traits which should have a particular value, we consider the optimal combination of traits to identify the occupiable regions of trait space which are most likely to lead to improved seed yield.

We therefore identified the points in the trait space which maximise the expected improvement in seed yield over the largest observed yield value in each of the Spring and Winter OSR panels. The identified points define “ideotypes” - hypothetical plants which are expected to produce high seed yield based on a model of the way that yield traits interact to produce seed yield. We only considered traits which affect yield directly (**Figure 3**). The other measured traits affect yield only through these, and so can in principle be ignored, assuming that the direct traits are controllable.

In Spring OSR, we found that when observed trait trade-offs were imposed as constraints, many of the observed plants were higher yielding than the best ideotypes identified. This implies that the high yielding regions of trait space are already well explored by the existing varieties, and it is not expected that other plants can be defined (at least in terms of the experimentally observed traits) which are expected to yield better under these growing conditions. Therefore, it is recommended that Spring OSR plants should produce 35-60 pods on the main branch, approximately 150-250 pods should be produced on secondary branches, seed number should be > 4000, seed weight per 10 pods on the main branch should be > 0.6g, TGW should be approximately 3g to 5g.

In Winter OSR, ideotypes were identified which are expected on average to perform better than all but the single best observed plant. This suggests that regions of the trait space exist which are likely be able to yield better than the existing varieties when grown under these conditions. The best identified ideotype has 90 pods on the main branch, and approximately 900 pods across 11 secondary branches. It produces just under 6000 seeds in total, with a relatively small TGW of 4g. Ovule area, and seed weight per 10 pods on the main branch is approximately average for winter varieties, implying a dichotomy between relatively large, high yielding pods on the main branch, and many small pods on the numerous secondary branches.

We constrained correlation relationships among yield traits to be the same as those observed in the experimental plants. This assumes that the observed relationships cannot be broken through selection, i.e. that they have unavoidable, physiological causes, (e.g. competition between traits for seed filling capacity) rather than correlated underlying genetic causes. If the observed correlations between traits can be broken, then these trait correlation constraints can be relaxed. We show that breaking these relationships is expected to result in large potential increases in seed yield, most simply through maintaining high seed size, and increasing seed number, as has been previously proposed (Labra et al., 2017). However, it is doubtful that these relationships can truly be relaxed, as it is generally considered that in OSR determination of seed number and size overlaps through competition for seed filling resources (Diepenbrock, 2000), accounting for at least part of the observed relationships.

Identified ideotypes are defined in terms of their desired trait values, however this does not address whether these ideotypes can be produced through selective breeding. High heritability is required for genetic selection. However, (as shown here), causal trait-trait relationships mean that often a large proportion of a trait’s heritability is not due to genetic variation in itself directly, but due to genetic variation in its ancestor traits. Consequently, if these ancestral traits are also under selection then the trait is in fact not as heritable as would be calculated naively, and selection upon it cannot be expected to be as effective. Therefore, given a list of multiple candidate traits for selection within a particular programme, residual heritability for each trait (conditioned upon the others) should be calculated in order to estimate whether they can be mutually selected upon successfully.

Yield is a complex trait, with many “yield traits” contributing to its value. Ideally, we would be able to learn seed yield directly as function of genotype, however (as we have shown through simulation) causal associations between traits lead to increasing genetic complexity, because genes which affect secondary yield traits also affect seed yield indirectly. Large statistical power is therefore required to identify the many of the genes which affect yield. Consequently, genes relevant for seed yield are often identified indirectly, through their association to traits which contribute to yield. This allows their association with yield to be identified through less powerful genetic association experiments than would be required to identify them directly. As we have seen, many other traits in OSR are also “complex”, as so can also benefit from a similar decomposition approach, and through careful control of contributing parent trait effects.

As discussed in the results, the potential cost versus benefit of correcting for causal trait-trait relationships depends on the relative frequency of causal trait relationships versus pleiotropic gene actions. This is currently unclear due to ambiguity between “true” pleiotropy, and mediated pleiotropy in many identified cross phenotype effects. The important distinction between these mechanisms will require careful experimentation to pull apart in future.

We have largely assumed that the trait-trait relationships inferred here are causal in nature, allowing a thought experiment into the consequences of causal relationships, and highlighting the importance of their consideration. As discussed in the introduction, this is reasonable in many cases, either due to developmental associations, physiological feedback regulating traits, or competition for resources between observed traits. However, the causality of trait-trait associations cannot be directly inferred from an observational study without making the unconfoundedness assumption – that all relevant variables are observed and included in the model. Here, this assumption may arguably be broken by some failure to include all relevant physiological traits, and by the omission of correlated genetic variation between varieties during the model inference process.

It is therefore not clear that all the links inferred in the model are necessarily directly causal. Linked traits may instead have a confounding common cause. Either an unobserved third trait, a shared pleiotropic gene, or a set of genes which act on the traits independently, but which are correlated in the population.

Whether a link is truly causal cannot be distinguished without perturbation experiments. These experiments generally have not been performed within the context of yield trait association studies (Aytaç & Kınacı, 2009; Chen et al., 2014; Engqvist & Becker, 1993; Golparvar, 2011; Ivanovska et al., 2007; A. Khan et al., 2000; F. A. Khan et al., 2006; Lu et al., 2011; Marjanović-jeromela et al., 2008; Marjanović-Jeromela et al., 2011; Özer & Oral, 1999; Rameeh, 2014; Sabaghnia et al., 2010; Tariq et al., 2020; Tunçtürk & Çiftçi, 2007).

The identified links therefore provide a set of hypothetical relationships consistent with the observed data, and prior information, and which should now be experimentally verified in an iterative process of model improvement.

Overall, we have shown the importance of accounting for trait-trait relationships in many facets of applied plant science, from identifying crop ideotypes, to estimating which traits can be selected for, to identifying gene-trait association. Accounting for these relationships will increasingly be both possible and important, due to the introduction of phenomics platforms.

All modelling here is based on the traits realised in individual plants, rather than the traits which can be expected from a particular genotype given a defined environment. We do not consider whether the identified crop ideotypes (defined in terms of desired trait values) can be reliably obtained in a given environment through genetic selection, let alone whether the expected region of trait space spanned by the progeny of a particular cross includes an ideotype plant. Coupling trait-trait modelling to genomic prediction models therefore has the potential to extend the utility of both approaches. Genomic prediction will allow an assessment of whether and how identified ideotypes could be produced, and trait relationship modelling can factorise the genomic yield prediction problem into a set of simpler subproblems. Together these quantitative modelling approaches may allow fully automated, optimal decision making for many aspects of plant breeding.

## Methods

### Plant growth conditions & trait measurement

The studied *B. napus* diversity set population consisted of 94 genotypes (Harper et al., 2012; Havlickova et al., 2017). The population was classified in 4 OSR groups, including Winter OSR (41 lines), Spring OSR (22 lines), Semi-winter OSR (8 lines) and Others (23 lines which included swede, kale, unspecified and fodder genotypes, **Supplemental Data Set 3**). For GWAS analysis, all lines were used, for all other analysis, only Spring and Winter OSR were used.

Plants were grown as described in (Siles et al., 2020) arranged in 2 glasshouses. Each glasshouse contained all 94 genotypes arranged in a 20x12 non-resolvable row-column design. All genotypes were replicated either 2 or 3 times per glasshouse to give a total of 5 replicates across both glasshouses. The design was generated in CycDesigN (CycDesigN 6.0, VSN International Ltd, Hertfordshire, UK).

27 traits were measured with either 3 or 5 biological replicates for microtraits and macrotraits, respectively (**Supplemental Table 2**). Phenotyping of these traits was as described in (Siles et al., 2020).

### Data pre-processing

The raw trait data was transformed to be normally distributed. The transformations applied to each trait are given in **Supplemental Table 3.** Missing values were imputed by predictive mean matching, using the “mice” (v3.11.0) R package (van Buuren & Groothuis-Oudshoorn, 2011) as detailed in **Supplemental Script “impute_data.R”**.

### Identification of trait relationship structure

Sequential Path Analysis models (Sabaghnia et al., 2010) were separately fit for Spring and Winter *B. napus* varieties. (**See Supplemental Script “id_trait_structure.R”)**. Based on previous biological knowledge of oilseed development, a hierarchy of permitted causal trait-trait relationships was defined, and a small set of links whitelisted for inclusion (see **Supplemental Data Set 4**). Within these constraints, relationships between traits were modelled by linear regression, to allow a rapid search of the permissible relationship space. Tabu search (implemented in the bnlearn R package, Scutari, 2010) was carried out to maximise the Bayesian Information Criterion (BIC) score of the trait-trait model. Bootstrap sampling of the data was used to estimate the probability of each inferred trait-trait link.

To reduce overfitting, the data was randomly split into five folds, and the modelling process carried out separately. The inferred models were averaged (following Scutari & Nagarajan, 2013) to only include links identified consistently.

### Predicting the consequence of modifying traits on seed yield

The inferred trait relationship structure (a directed acyclic graph, DAG) was used to define non-linear regression models of each observed trait. The predictor variables for each trait are the parent nodes of that trait in the DAG. Modelling was carried out in the Stan language (Stan Development Team, 2020).

To predict the effects of singly modifying each trait between its extreme observed values, it was set to a modified value, and the estimated values of its direct children sampled. These predictions were then used to estimate the values of the modified-trait’s grandchildren and so on, respecting the dependencies identified in the DAG, until all descendants of the modified node were predicted. Traits which were not descendants of the modified trait were held to their observed *in planta* values.

For each modelled trait, we used a Gaussian Process (GP) prior with an Automatic Relevance Determination kernel

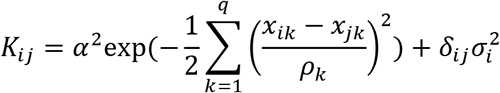

where *q* is the number of parent traits, and *δ_ij_* is Kronecker’s delta (1 if *i* = *j* and 0 otherwise). Hyperparameters *α ρ*, *σ* were estimated through regularised maximum marginal likelihood with priors *α* ∼ *N*(0,1), *ρ_k_* ∼ *IG*(5,5), *σ*_i_∼ *N*(0,1). Predicted trait values were estimated by sampling from the posterior distribution to allow uncertainty in predictions to propagate through the DAG structure.

The predicted effect of modifying each trait in Spring and Winter OSR were averaged over the 5 candidate trait-relationship structures identified. Code used to estimate the effects of modifying traits is provided in **Supplemental Script “trait_modification_prediction.R”**.

### Bayesian optimisation for ideotype identification

To identify optimal crop ideotypes for maximum seed yield and propose other promising regions of trait space for exploration, we used a Bayesian Optimisation framework. As the “surrogate model” we used the empirical GP models for predicted seed yield conditioned on the observed values of the parent traits of seed yield in either Spring or Winter OSR DAGs.

As the “acquisition function” for maximisation we used Expected Improvement (EI, Mockus et al., 1978) defined as

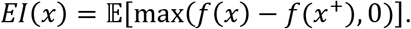

Where *f(x)* is the predicted value of seed yield at *x*, and *x*^+^ is the observed point in trait-space with the maximum seed yield. EI thus allows us to balance exploitation of regions with high predicted seed yield, with exploration of regions with high posterior uncertainty.

Proposed trait values were constrained to be between the maximum and minimum observed values.

Because generating crosses is costly, but parallelizable, it is desirable to be able to propose multiple points for exploration given the current data. We used the “Constant Liar” heuristic strategy proposed in Ginsbourger et al., (2010), with L equal to the maximum observed seed yield in order to obtain an approximately q-EI optimal design for *q* proposed sampling points.

Relationships exist between the yield traits exist in the observed data, and so it is unlikely that all points of trait-space can be occupied by a real plant. To incorporate correlation between the parent traits as constraints in the optimisation of proposed points, we followed the approach proposed in MacGregor, (2004). An independent (correlation free) basis for the trait-space was identified by PCA of the parent traits of seed yield. PCs sufficient to explain >95% of variation were used as predictors, and Bayesian optimisation was carried out using this new independent basis as described above. Predictions in the PC basis were then transformed back to the observed trait basis for reporting.

Code used for crop ideotype identification is provided in **Supplemental Script “bayesian_optimisation.R**”.

### Consequences of trait relationships for heritability and genetic variant association

See **Supplemental Methods** for details of the regression analyses carried out in order to ascertain to consequences of identified trait-trait relationships on heritability estimates, and GWAS analysis.

## Author contributions

LS carried out all experimental work. AC developed the computational approaches, carried out the majority of data analysis, prepared the figures and the first draft. LS, AC, RJM, SK wrote the manuscript with contributions from PE. SK and RJM supervised the project.

## Acknowledgements

The authors would like to thank Dr Christina Sanchis-Gritsch (Rothamsted Research, UK), and Hannah Walpole (Rothamsted Research, UK) for their assistance in data collection and analyses. We thank Dr Kirsty Hassall (Rothamsted Research, UK) for her help in statistical processing of the collected data.

## Supplemental Figure legends

**Supplemental Figure 1: Traits may be correlated to seed yield without a causal relationship between them** Predicted trait values (y-axis) as traits a singly varied (x-axis). Median prediction shown with a heavy line, confidence quantiles are shown with 10% intervals.

The predicted effect of modifying **a)** silique beak length, **b)** pod length on other yield traits and seed yield in Winter OSR. Changing beak length is not expected to much affect yield. Modifying pod length is expected to strongly affect both beak length and seed yield. Consequently beak length and seed yield are correlated. **c)** a trade-off exists between seed number and thousand grain weight (TGW) in both Spring and Winter OSR.

**Supplemental Figure 2: Relevance of directly and indirectly connected yield traits for ideotype design. a)** Example trait-trait model, in which seed yield is directly affected by a secondary yield trait, and indirectly affected by two tertiary yield traits. Seed yield is independent of indirectly connected traits, conditional upon the traits which link to it directly. **b)** Consequently, the dimensionality of the yield surface can be reduced to only include those secondary traits directly connected to yield. Therefore, if breeding material with appropriate secondary trait values exist, then indirectly connected traits can be ignored, and the optimal crosses identified through consideration of secondary yield traits alone. **c)** Only if parents with appropriate secondary traits do not exist is it necessary to consider indirectly linked traits. In this case, the directly connected secondary yield traits can be considered in terms of the tertiary yield traits which cause them in order to ascertain crosses which are likely to generate progeny with the desired secondary trait value, and therefore seed yield.

**Supplemental Figure 3: OSR ideotypes identified ignoring empirically observed relationships between traits**

Facets show traits which affect seed yield directly in the **a)** Spring and **b)** Winter OSR panels.

Coloured points show mean predicted seed yield for identified hypothetical ideotype plants with the indicated yield trait values. Their order reflects their probability of having greater seed yield than the best experimentally observed plant, with 1 having the best chance, and 10 the worst of the calculated ideotypes. These points are identified through Bayesian optimisation as the unobserved points with the best chance of exceeding the maximum observed seed yield in each panel. Black points show seed yield and yield traits for experimentally observed plants. Thin black line shows the maximum observed seed yield produced by any of the plants in the Spring or Winter panels, thick black line shows mean observed seed yield.

The yield traits exhibit correlations in the observed data, but it is not clear whether this is an inevitable consequence of trade-offs among them, or whether these correlations exist due to related genotypes, and could ultimately be at least partially overcome. If expected improvement in seed yield is maximised without the constraint of respecting observed correlations between yield traits, then it is clear that more optimal regions of trait space exist than the points occupied by observed plants in both spring and winter OSR. As might be expected, this is largely via breaking the negative trade-off between seed size, and seed number in both Spring and Winter OSR.

**Supplemental Figure 4: Many yield traits can be well predicted from their parent traits**. Plots show the mean predicted trait values vs observed trait values for each plant in **a)** Spring OSR, **b)** Winter OSR. Variance in the trait explained by the parents is given above each plot. It can be seen that variation in many traits can be well explained by their parent traits, indicating that this variation may not be due to direct genetic or stochastic variation in the trait itself, but instead due to variation in its parent traits.

**Supplemental Figure 5: Simulation shows that unless accounted for, trait-trait relationships reduce power to detect gene associations as well as the misleading identification of indirectly associated genes.**

**a)** Model from which simulated data were generated for each plant independently (j). Parent-trait (p) is the weighted sum of “parent SNPs” (s_1_, … s_n_) which affect it directly. Child-trait (c) is the weighted sum of “child SNPs” (r_1_, …, r_m_), as well as the parent trait. So, it is affected directly by the child SNPs, and indirectly by the parent SNPs. Parent SNPs will therefore exhibit mediated pleiotropy. Noise (ϵ^p^, ϵ^c^) was added to both traits. “Non-causal SNPs” (q_1_, …, q_k_) do not affect either trait. All SNPs were independently sampled from a Bernoulli(0.5) distribution.

**b)** The number of “parent SNPs”, “child SNPs” and “non-causal SNPs” statistically associated with variation in the child-trait, using either observations of the child trait directly (red), or correcting for trait-trait relationships, by using the residuals of a model in which child-trait was predicted by parent-trait (blue). (See methods section for details of associated SNP inference). In all tested cases, (except when c is independent of p, *γ* = 0), the power to detect child SNPs is greater when the effect of p on c is controlled for. The greater the value of *γ*, the bigger the difference. When *γ* is large relative to *σ*, parent SNPs are identified rather than child SNPs due their indirect effect (for example when *γ* = 3, *σ* = 0.1). When *γ* is similar to *σ*, association using the direct observations of c was less able to detect any directly or indirectly causal SNPs (for example when *γ* = 1, *σ* = 0.5). Neither method was more associated with spurious identification of non-causal SNPs.

**Supplemental Figure 6: Correlations between traits and yield vary based on environment, and genotypes considered.**

The measures of yield shown are “SeedYield” (weight of seed produced per plant), or “SeedYield (kg / hectare)”. Lu et al, Jeromela et al, Aytac & Kinaci 2003 & 2004 use Winter OSR panels. Chen et al use Spring and Winter OSR, the remainder use Spring OSR panels. Years indicate repeated trials in the same study, with the exception of Khan 2000, Khan 2006 which are separate studies. Sabaghnia et al., and Ivanovska et al., alter environmental conditions within a study, either experimentally or through trial location. Referenced studies are (Aytaç & Kınacı, 2009; Chen et al., 2014; Golparvar, 2011; Ivanovska et al., 2007; A. Khan et al., 2000; F. A. Khan et al., 2006; Lu et al., 2011; Marjanović-jeromela et al., 2008; Özer & Oral, 1999; Rameeh, 2014; Sabaghnia et al., 2010; Tariq et al., 2020; Tunçtürk & Çiftçi, 2007).

**Supplemental Figure 7: QQ-plots for models to associate SNP data with the number of pods on the main inflorescence (NumberPods M), and seed oil content (OilContent)**. QQ plots are shown for models associating SNPs to the trait directly (raw), and to the residuals of models in which the trait is predicted from its parent traits.GLM-Q, GLM-PCA5d, GLM-PCA10d are general linear models (GLM), which use the SNP data, as well as either the population structure matrix (Q), first five (PCA5d), or first ten principle components (PCA10d) of the SNP matrix. MLM-K, MLM-Q+K, MLM-PCA5d+K are mixed linear models (MLM) which use the SNP data, as well as the Q-matrix, PCA components, or kinship matrix (K). A Bayesian-information and Linkage-disequilibrium Iteratively Nested Keyway (BLINK) model was fit using the GAPIT package implementation. Based on these plots, GLM-PCA5d was used for NumberPods M, and GLM-PCA10d was used for OilContent.

## Supplemental table legends

**Supplemental Table 1:** Estimated genetic control of phenotypic traits. Statistical significance of genotype effect on measured traits estimated by one way ANOVA. Reported p-values were adjusted for multiple hypothesis testing by Benjamini-Hochberg method. Broad sense heritability 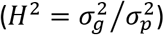 was estimated from the mean squares components of ANOVA, following (Singh et al., 1993), using either normalised observed trait values, or model residuals (see methods). Dashes indicate that the trait is not modelled as having any parent traits, and so residual values are the same as for raw values.

**Supplemental Table 2:** List of macrotrait (n=5) and microtrait (n=3) names and abbreviations measured in the diversity set population.

**Supplemental Table 3:** List of transformations applied to normalise trait distributions.

## Supplemental data set legends

**Supplemental Data Set 1: details of SNPs associated with Number of Pods on the main inflorescence trait.**

**Supplemental Data Set 2: details of SNPs associated with the oil content trait.**

**Supplemental Data Set 3: List of 94 genotypes included in the diversity set population.** The ASSYST code, genotype names, crop type description and the 4 oilseed rape groups are presented.

**Supplemental Data Set 4: Trait-trait relationship constraints for sequential path analysis**

